# Analysis of viromes and microbiomes from pig fecal samples reveals that phages and prophages are not vectors of antibiotic resistance genes

**DOI:** 10.1101/2021.05.20.444921

**Authors:** Maud Billaud, Quentin Lamy-Besnier, Julien Lossouarn, Elisabeth Moncaut, Moira B. Dion, Sylvain Moineau, Fatoumata Traoré, Emmanuelle Le Chatelier, Catherine Denis, Jordi Estelle, Caroline Achard, Olivier Zemb, Marie-Agnès Petit

## Abstract

Understanding the transmission of antibiotic resistance genes (ARGs) is critical for human health. For this, it is necessary to identify which type of mobile genetic elements is able to spread them from animal reservoirs into human pathogens. Previous research suggests that in pig feces, ARGs may be encoded by bacteriophages. However, convincing proof for phage-encoded ARGs in pig viromes is still lacking, because of bacterial DNA contaminating issues. We collected 14 pig fecal samples and performed deep sequencing on both highly purified viral fractions and total microbiota, in order to investigate phage and prophage-encoded ARGs. We show that ARGs are absent from the genomes of active, virion-forming phages (below 0.02% of viral contigs from viromes), but present in three prophages, representing 0.02% of the viral contigs identified in the microbial dataset. However, the corresponding phages were not detected in the viromes, and their genetic maps suggest they might be defective. Furthermore, our dataset allows for the first time a comprehensive view of the interplay between prophages and viral particles.

## Introduction

The spread of antibiotic resistance genes (ARGs) is a major health concern, and powerful sequencing techniques have made it possible to closely examine the contribution of mobile genetic elements to their spread. The main mobile vectors that are responsible for spreading ARGs are conjugative plasmids and integrative conjugative elements (ICEs, also known as conjugative transposons). In both cases, ARG spread occurs through conjugation, a process that involves the formation of a conjugative pilus, the contact between a donor and a recipient bacterium, and the transfer of the genetic element. ARG are frequently found in pig fecal samples^1^, and their associated mobile genetic elements start to be well documented. For instance, analyses were conducted on a set of metagenomics contigs that were cloned on bacterial artificial chromosomes. Eleven of them were conferring resistance to tetracycline to *Escherichia coli*, and nine of the *tet* genes were present on mobile genetic elements (plasmids or ICEs)^2^.

Interestingly, bacteriophages, another category of mobile genetic elements, apparently rarely encode ARGs^3^. Phages can usually be divided into two large groups based on their lifestyle. Virulent phages inject their genetic material into bacteria, replicate and lyse their host cells, thereby releasing several new virions at the end of this lytic cycle. Temperate phages are able to alternate between the lytic cycle and a dormant stage where they maintain their genetic material within the bacterial genome as prophages (following an integration step or as plasmids). While dormant, prophages may still express a few of their genes, including some are beneficial to their host, such as *bor*, which is involved in resistance to serum complement killing^4^, and the iron transporter *sitABCD* genes^5^. In theory, a temperate phage and its expression profile in a prophage state would offer an efficient process for the dissemination of ARGs. So, why have they so rarely been detected?

Initially, their absence was noted on the basis of single phage biology methods, such as phage culturing and sequencing, and therefore the investigations lacked breadth. On the other hand, qPCR studies have reported the presence of ARGs in virus-enriched environmental samples from natural waters^6^, wastewater plants^7^, or human/animal fecal samples^8,9^. However, virome preparations are often highly contaminated by bacterial DNA^10,11^, making it difficult to discern whether the ARG originates from viral or bacterial DNA using qPCR alone. Another confounding factor is the process of generalized transduction. During this process, instead of packaging their own genome, some phage particles transport bacterial DNA (sometimes at high frequencies^12^).

Shotgun metagenomics studies of enriched viral fractions (viromes) and improved programs for assembling short reads into quality contigs can now help determine whether ARGs are found in phage genomes. ARGs were recently investigated in the virome from wastewater samples, and they were found to be scarce compared to other genetic elements^13,14^. However, several reports have raised a concern with pig fecal samples, which appeared to be rich in phage encoded ARG^8,15,16^.

Another approach for investigating the ARG content of phages is to study prophage regions of completely assembled bacterial genomes. Recent reports on pathogenic bacterial species such as *Acinetobacter baumanii* or *Streptococcus suis* found prophage regions with ARGs^17–19^. In these *in silico* studies, the difficulty lies in the ability to discern whether such prophages are defective or still functional and able to, for instance, complete lytic cycles, form viral particles, and therefore spread ARG in bacterial populations.

Here, with respect to the ARG content in pig fecal viromes, we elected to extract and analyse new samples, because the presence of bacterial DNA contamination was either clearly mentioned in previous reports^16^, or it was poorly documented, by the absence of detection of a DNA band after PCR with 16S universal primers. Moreover, we sequenced in parallel the purified virome and the total microbiota of the same samples to have access to the prophage content and be able to test whether they were active (i.e. present also in the virome fraction). Our study presents the results of these analyses performed using 14 pig fecal samples collected from farms with varying degrees of antibiotic usage and reports the presence and absence of ARG among microbiota and viral particles, respectively.

## Materials and Methods

### Sampling

Fourteen farms in Brittany (France) were selected during year 2015, for pig feces sampling based on their various veterinary practices and more precisely, on their antibiotic treatment protocols. The antibiotic treatments for the fourteen animals used in our study are shown Suppl. Table 1. Since piglets are more heavily treated with antibiotics, five of the 14 samples were collected from piglets rather than adults.

### Virome DNA sample preparation and sequencing

Viral particles were enriched from fecal samples following the route 5 protocol described in Castro-Mejia et al. ^20^ but with minor modifications. Two grams of frozen fecal sample were resuspended into 50 mL of PBS adjusted to 0.2 M NaCl, homogenized and agitated overnight at 4°C on a rolling device. After a prefiltration step on glass-fiber filters, samples were centrifuged at 5000 g 30 minutes at 4°C, and bacterial pellets were discarded. This step was repeated until supernatants were clear (up to 3 times). Five grams of polyethylene glycol 8000 was then dissolved into the phage supernatants, and phages were left to precipitate overnight at 4°C. The precipitate was then centrifuged for 10 minutes at 15,000 g, after careful removal of PEG, and resuspended into 1 mL of SM buffer (50 mM Tris pH 7.5, 100 mM NaCl, 10 mM Mg-SO4). A iodixanol-SM step gradient was then formed in a 12-mL ultracentrifugation tube (Beckman, ultraclear tubes), by applying 6 mL of a solution at 20% iodixanol-SM, and then adding slowly with a Pasteur pipette, below this volume, 4.5 mL of a 45% iodixanol-SM solution. Finally, the phage sample (1.5 mL) was placed on top of the tube. The sample was ultracentrifuged for 5 h at 40 000 g, at 10°C. Most of the 14 gradients contained two visible bands. The lower band was the viral band, which was collected from the bottom of the tube and dialyzed against SM buffer (4 h) in two successive 1-L volumes (final volumes 3-5 ml after dialysis). Samples (0.9 mL) were then treated with DNAseI (2.5 U, 1h30 with Turbo DnaseI, Ambion) to remove residual free DNA.

To check for bacterial DNA contamination of the virome preparations prior to DNA extraction, qPCR was performed with universal 16S primers and compared to an *E. coli* total DNA standard. Total nanoparticle concentrations were also estimated with a Videodrop apparatus (Myriade), an apparatus that detects nanoparticles using light interferometry^21^. Most samples had less than 10 ng/mL of bacterial DNA for every 10^10^ nanoparticles and were not further treated prior to gradient loading. Filtration (0.2 μM pore size, polyethersulfone filters) was performed on four samples (VP4, VP5, VP15 and VP17) that were considered too heavily contaminated with bacteria. For viral DNA extraction, 1 mL of viral sample was added to a 1-mL solution of 50% phenol/50% chloroform-iso-amyl alcohol (24:1), manually mixed for 1 min, and centrifuged for 20 min at 12 000 g. The soluble phase was recovered and treated a second time with phenol-chloroform. The soluble phase was then added to an equal volume of chloroform-isoamyl alcohol (24:1), emulsified and centrifuged for 5 min at 12 000 g. DNA from the soluble phase was finally precipitated by adding sodium acetate (0.3 M), 1 μg of glycogen as the DNA carrier, and 0.6 volume of isopropanol. The DNA pellet was collected by centrifugation, washed with 70% ethanol, dried, and resuspended in 11 μL of 10 mM Tris pH 8. DNA was quantified using Qubit. Due to the insufficient total amount of DNA obtained (20 to 250 ng out of the 600 ng required), a multiple displacement amplification step (MDA, with kit Genomiphi2) was performed on 1 μL of each sample (90 min at 37°C), after microdialysis (0.02 μM VWPS filters, Millipore) to remove potential inhibitors. This step allowed for the inclusion of ssDNA viruses in the analysis. However, RNA viruses (which are the less abundant among known bacterial viruses) were excluded. Libraries were prepared using the Truseq kit and sequencing was performed on an Illumina Hiseq platform (2 x 150 nt, pair end), with an average depth of 50 million total reads per sample.

### Virome read cleaning, assembly and contig treatments to constitute the VP reference dataset

Reads from each sample were trimmed using Trimmomatic^22^ (ILLUMINACLIP:TruSeq3-PE.fa:2:30:10 LEADING:3 TRAILING:3 SLIDINGWINDOW:4:20 MINLEN:100), and dereplicated with the fastx-unique tool from the free version of the Usearch9 suite (R. Edgar, http://drive5.com/usearch). Reads were then paired again using the fastq_pair program (Edwards Lab). Assemblies were run separately on each sample using SPAdes (Bankevich et al., 2012) (version 3.13, parameters: --only-assembler -k 21,33,55,77,99,127) to include both paired and unique reads. The meta option in SPAdes was disregarded to improve contig lengths. A clustering step of viral contigs from all samples was performed at 95% nt identity using CDhit (psi-cd-hit.pl -c 0.95 -G 0 -g 1 -aS 0.9 -prog blastn) and only contigs that were equal to or above 2 kb were retained for further analysis. The final viral dataset contained 7755 virome-de-porc (VP) contigs (out of the original 8090 contigs prior to clustering).

To examine the assembly completeness of the VP contigs dataset, dereplicated reads were mapped back on the 7755 contigs using the Bowtie2^23^ with default parameters (allows multiple matches, with random affiliation). On average, 75% of total virome reads were ampped back on the VP collection. To generate abundance matrices, mapping result were treated with tools from the Samtool suite^24^ (view, sort, index, idxstat), and the number of reads found on each contig was finally converted into reads per length of the contig (in kb) per million (RPKM). The VP matrix contained high RPKM values, ranging from 5 to 91 000.

### Microbiota DNA sample preparation and sequencing

Total microbiota DNA extraction was performed according to protocol H of the international standards for human fecal samples (IHMS, http://microbiome-standards.org/index.php?id=Sop&num=007), starting from 0.2 g of frozen fecal sample. In short, purification steps involved guanidine thiocyanate/SDS sarkosyl treatment for 1h at 70°C, followed by bead beating to break open all bacteria, then repeated polyvinylpolypyrrolidone applications to remove polyphenols. DNA was then precipitated with isopropanol, washed and dried. This protocol should also break open the viral capsids, thanks to the initial guanidine thiocyanate protein denaturation step, although the question has never been specifically addressed, to our knowledge. However, the absence of any step to convert ssDNA into dsDNA prevented sequencing of ssDNA viruses. Libraries were prepared using the Truseq kit and sequencing was performed on an Illumina Hiseq platform (2×150 nt, pair end), with an average depth of 61 million total reads per sample.

### Microbiota read cleaning, assembly and contig treatments to constitute the P reference dataset

The same steps applied for viromes were followed for total microbiota, with some simplifications. Briefly, reads of each sample were trimmed with Trimmomatic (same parameters as virome reads, except MINLEN=125), and then directly assembled with SPAdes (same options as for viromes). A clustering step of of each individual microbial sample was performed at 95% nt identity with CDhit, and only contigs of a size equal to or above 2 kb were retained for further analysis. The microbial dataset comprised 220,000 pig (P) contigs (from 271,163 contigs prior clustering).

To examine the assembly completeness, reads were mapped back onto the P contig dataset using Bowtie2 on default parameters. On average, 69% of microbial reads were mapped back on the P contigs collection. To generate abundance matrices, mapping results were treated with tools from the Samtools suite, and the number of reads found on each contig was finally converted into RPKM. The P contigs matrix contained RPKM values that were 10-times lower than the VP matrix, ranging from 0.71 to 6890.

To estimate the main phyla present in the microbiome samples, a taxonomic affiliation was determined using Kaiju ^25^ for the top 10 585 P contigs with a global abundance above 10 RPKM. To sum up contig abundances by phyla, those which might belong to the same microbial species were binned into clusters based on kmer composition and co-occurrence with VAMB ^26^. A single contig per VAMB cluster was conserved in the abundance matrix, which ended up containing 5862 contigs (~species). Moreover, in the context of ARG analyses, Kaiju-based taxonomic assignment of interesting contigs was complemented with a BLASTn against the nt database (retaining the best hit with E-value below 10^−50^, when present).

### Bacterial contamination level of the viral-enriched DNA

The amount of bacterial DNA contamination of the virome reads was estimated using two methods. First, reads were mapped onto the 16S SILVA collection^27^ with Bowtie2. All samples had less than 2 x 10^−4^ reads homologous to 16S, which is a cut-off for viromes with satisfactory-level purity^11^. The maximal 16S read ratio was 3 x 10^−5^ in sample VP2 (see Suppl. Table 2). Second, reads were submitted to the viromeQC program^10^, which estimates a viral enrichment yield. This is done by comparing the frequency of reads matching with 16S, 23S and ribosomal protein encoding genes, to that expected for global microbial metagenomics samples. Enrichment was 100-fold, except for 3 samples (lowest value 7-fold, sample VP2, Suppl. Table 2). ViromeQC results matched well with the 16S method, and showed that the pig viromes were essentially viral in content.

### Analysis and labeling of the bona fide viral contigs within the VP dataset

From the 7755 VP dataset, the 2480 contigs with the highest RPKM (above 100) were first analyzed. Several programs and methods were applied to identify non viral contigs (bacteria and plasmids, see suppl. Figure 2). Virsorter ^28^ recognized 23% (568 contigs) of these contigs as viral (all categories were accepted, cat1 and cat2 were vastly dominant). The Inovirus detector ^29^ was then run on the remaining contigs and 143 inoviruses (putative or confirmed, most were circular). To add contigs corresponding to phages recently added into public databases, MegaBLAST was run against the nt database of NCBI (April 2020, E-val below 10^−16^). As bacterial hits could also correspond to prophages, all BLASTn hits, as well as the remaining dark matter were taken for a search centered on proteins, making use of PHROGs, a database of phage proteins (Terzian et al., submitted). For this, an HMMER hmmscan ^30^ was used for proteins from all remaining contigs against PHROG profiles was run, and all contigs with at least 5 orfs and 35% of them homologous to a PHROG family (E-value below 10^−12^) were counted as phages. In total, this BLASTn + PHROGs step allowed for the recognition of 966 additional contigs from the following origins: 654 phage, 232 eukaryotic viral (collectively designated below as “viruses” for short, while “phages” is kept for bacterial viruses), 72 bacterial and 8 plasmid fragments. Next, the VIBRANT classifier ^31^ was used on the remaining uncharacterized contigs, leading to additional 325 phages and 29 viruses. In the end, only 449 contigs of the 2480 (18%) remained uncharacterized.

All previously accepted phages were verified using VIBRANT and revealed a high degree of overlap for all *Caudovirales* and *Petitvirales* genomes (92.5% and 99% were detected by VIBRANT, respectively). VIBRANT was also successful at detecting eukaryotic viruses, as 79% of them were detected. VIBRANT was less successful at detecting inoviruses (35% detected). We then used VIBRANT and the Inovirus detector on the remaining 5275 low-abundance contigs and obtained an additional 3654 viral and 201 inovirus contigs. In total, 5806 contigs were recognized as viral, and among those treated by VIBRANT (contigs with ≥ 4 ORFs), 77.3 % were viral.

### Contig annotations and ARG search

VP and P contigs were annotated using Prokka ^32^ (version 1.13 with --addgenes and –rnamer options). Following annotation, three ARG searches were compared for the VP contigs: (i) hmmscan against ResFam profiles (^33^ –cut_ga option), as in our previous report ^3^; (ii) BLASTp against the MUSTARD database ^34^ (≥ 95% id and ≥ 90% alignment cut-offs), built in part with ARG genes found in the human gut; (iii) BLASTn against the version 4.0 release of Resfinder ^35^, with parameters ≥ 90% identity, ≥ 60% coverage. For microbial P contigs, only the Resfam and Resfinder searches were performed. A deeper analysis was then conducted on the most prevalent contigs. BLASTn search against nt (as of November 2020) with contig P4_NODE_19877 (2256 bp, contains a *tetR* gene), indicated that among the top 176 BLASTn hits (≥ 99% identity, 100% coverage), 41 were *Clostridiales*, and 11 *Bifidobacteriaceae* complete genomes. In the vicinity of the *tet* gene, genes annotated as *traG, mobC* and relaxase were found, suggesting the presence of an ICE.

### Elements of VP contig classification

#### single-stranded (ss) DNA viruses

A total of 2807 ssDNA viruses were recognized. *Petitvirales* were recognized either because they were specifically mentioned (*“non-Caudovirales* gene enrichment” score) in the Virsorter output, or by searching for terms ‘VP1’, ‘VP4’, and ‘Pilot’ in the annotated contigs of the VIBRANT output. Most ssDNA eukaryotic viruses were detected thanks to Megablast searches against the viral section of nt, and therefore directly affiliated. A ‘REP_TLCV’ (rep gene of tomato-leaf curl virus) annotation in VIBRANT files allowed to detect some additional eukaryotic viruses. *Tubulavirales* were detected thanks to Inovirus-detector ^29^. Most (272 of 344 total inoviruses) were circular, and these were analyzed into more details. A vCONTACT2 clustering revealed 50 clusters with a quality above 0.5, involving altogether 65% (179 phages) of all input genomes. Pairwise ANI (average nt identity) were also computed for all genomes sharing the same *gpI* gene, a gene typical of filamentous phages and needed for virion morphogenesis. Species membership was defined for pairs with ANI above 0.95, and genus boundary ANI > 0.75. A summary of the 272 circular genome properties is listed in Suppl. Table 3.

#### double-stranded (ds) DNA viruses

All viral contigs not recognized as ssDNA viruses were placed abruptly into the *Caudovirales* category (2999 in total). These were clustered with vCONTACT2 ^36^, together with the 2191 reference phages from Viral Refseq v88, and 23 additional intestinal phagessuch as LAK phages^37^, phages infecting *Faecalibacterium prausnitzii*^38^ and *Roseburia intestinalis*^39^. For 1675 of the 2999 VP contigs, a cluster was formed (478 clusters altogether), but only 18 of them contained reference phage genomes. All clusters of a quality above 0.5 (as indicated in the ‘quality’ column of the clusters output file of vCONTACT2) were considered further, and the top 7 are listed in Suppl. Table 4. These vCONTACT2 clusters were used to coalesce the most abundant *Caudovirales* contigs.

#### Huge phages

VP homologs to the 361 huge phage genomes^40^ were searched for by two approaches: first, the whole VP dataset was compared with BLASTn (E-val < 10^−5^) to the huge phages. Very few hits were found (30) and none with a query coverage above 14%. Then a more sensitive search was done with tBLASTx on a subset of large virome VP contigs and microbiota P contigs. These included the 21 VP contigs of a size above 150 kb, a 381 kb P contig considered viral due to its homology to a VP contig by BLASTn, and 5 VAMB bins having both a Kaiju annotation as viral and a cumulated size above 180 kb. These contigs were compared to the 35 closed_circular_curated huge phages^40^ and the 15 complete *Prevotella* megaphages^37^ with cutoffs placed at 30% query cover, 65% (amino-acid) identity and E-value 10^−3^.

Phage genome annotations were performed by running a BLASTp of ORFs predicted with Prokka against the NR database at NCBI (E-val < 10^−5^), and if needed HHpred against PDB (P > 98%) using the online Tübingen Toolkit Webserver^41^, and alignments maps were done with Easyfig^42^.

### Partitioning active phage versus inactive prophage contigs in the microbial P dataset, among contigs detected as viral with VIBRANT

A schematic summarizing the classification of active phage and inactive prophage contigs is shown in Suppl. Figure 3. Among the 220 000 P contigs, the viral ones were searched with VIBRANT. This gave 16 612 positive contigs. To complement this collection, 328 contigs matching by BLASTn to a VP contig (> 95% nt identity) or affiliated by Kaiju as viral, were added (total 16 940 viral P contigs). We reasoned that this set of viral contigs represented two very different situations, either virulent or temperate phages replicating and forming virions (category “active phage”), or dormant prophages that do not manifest viral activity (ie no capacity to form a free virion, at least in the ecosystem studied, category “prophage”). We took advantage of the virome reads to partition this set into either “active phage” if the viral P contig was matched by virome reads, or “prophage” if not, as described below.

We were also interested in the overlap between VP contigs and these viral P contigs. In a first step, therefore, the virome VP contigs were placed together with microbiome viral P contigs and dereplicated (95% identity, 90% coverage), generating a set of 23 181 non redundant viral contigs. Among them, 6658 were found only in the VP set (54 Mb cumulative length). There were 660 contigs in the “overlap” category, where contigs are representing both viral P and VP segments, with in some cases many small P contigs matching a large VP contig, and also the reverse situation, indicating that viral assembly was incomplete in both cases, unsurprisingly (24 Mb). The last 15 873 contigs were found only in the P assembly (178 Mb).

In a second step, virome reads were mapped onto the complete collection of 23 181 non redundant viral contigs. We reasoned that the modest overlap of the VP contigs with P ones (only 14% of VP have a match to P) could be the result of partial assembly, so that parts of the same phage may be assembled in each set. Therefore, it was expected that some of the P contigs having no overlap with any VP contig might still be corresponding to active phages. In this mapping, the VP contig with lowest cumulative abundance across the 14 samples had a RPKM value of 2.6. We then considered as “active viral contigs” all P contigs with global RPKM abundance above 2.6. This added 3049 P contigs as active viral contigs (39 Mb). In total, of the 16 940 starting viral P set, 4021 contigs (23.7%) belonged to the active phage category (corresponding to these 3049, together with 972 P contigs present or included into VP contigs of the “overlap” set), while the rest was considered inactive prophages.

### Host predictions for interesting phage clades

For the 7755 VP contigs, host prediction was first attempted by searching for matching CRISPR spacers, using the recent spacer database from Dion et al.^43^ and the default parameters (a maximum of two mismatches allowed). A host was predicted for 469 of these contigs (6% of total contigs). Next, a set of spacers was extracted from our P contigs (of all sizes, including those below 2 kb). For these spacers, CRISPR arrays were first identified using CRISPRDetect v2.2^44^. Spacers were then extracted from the CRISPRDetect GFF output file along with the following metadata: contig ID, start and end positions on the contig, length, sequence, orientation and position inside the locus. Spacers were assigned a unique ID and a custom BLAST database was generated (n = 14 941 spacers). Next, BLASTn v2.9.0 was used to perform pairwise alignments between the P contig spacers and the selected viral contigs, applying the same filters as previously described^43^. A total of 1075 (14%) VP contigs matched the in-house spacer collection. For 148 of the P contigs containing such matching spacers, a taxonomical affiliation could be proposed, based on the Kaiju indication followed by a BLASTn search (E-value < 10^−16^) against the Bacteria section of the nt database. As expected, affiliation was more successful using the in-house spacer collection, although in most cases, taxonomical affiliation of these in-house P contigs was unsuccessful (many very short contigs).

For the two clades of phages mostly focused in our study, 272 inoviruses and 94 Oengus-like phages, 15 and three hosts were predicted by this first method, respectively. Three additional tests were performed on this subset: (1) submission to the spacer collection at JGI (https://img.jgi.doe.gov/cgi-bin/vr/main.cgi) and applying the two-mismatch cutoff as above, (2) BLASTn search against NCBI nt (E-value < 10^−4^ and coverage > 10%), and (3) WiSH^45^. WiSH computes conditional probabilities of all 9-mers in bacterial genomes of the database (2155 bacterial genomes meaningful for the ecosystem were taken) and compares them to that of the query phage. The first output is the probability (expressed as log-likelihood) that the given phage will infect each of the bacteria in this collection. The closer the log-likelihood is to 0, the more likely the host is correct. A log-likelihood is also calculated for the phages that do not infect the bacteria of this same collection. By comparing these two log-likelihoods, a second statistic (a p-value) gives the confidence of the WiSH prediction. Four inoviruses and four phages of the Oengus cluster with known hosts were tested as controls. Only the five best log-likelihood with a p-value inferior or equal to 0.05 for each phage contig were considered as putative hosts. Next, a filter was applied on the log-likelihood values, which had to be below those obtained for the control phages. Finally, the bacteriophages with 4 to 5 predictions from the same taxonomic order were considered sure and kept.

### Data availability

All sequencing data are being submitted to ENA, experiment number ERX5476850.

## Results

### Among most abundant virome contigs, only 18% correspond to viral dark matter

Viral fractions of the 14 pig fecal samples were purified, separated on gradients, their ss- and ds-DNA extracted, amplified by multiple displacement amplification (MDA), and sequenced, as described in Materials and Methods. Reads were assembled into contigs, those of a size above 2 kb were retained, and clustered at the species level (>95% nt identity). Viral reads were mapped back onto the 7755 contigs obtained, and relative abundance of these contigs were computed to generate the so-called “VP” (for ‘virome-de-porc’) matrix.

The 2480 contigs with total RPKM values above 100 were then selected for in-depth viral sequence identification (see Materials and Methods and Suppl. Fig. 1A). Overall, 82% of all abundant contigs (2031) could be affiliated, therefore reducing unclassified contigs (the so-called “viral dark matter”) to only 18% of the sub-set of abundant VP contigs (Suppl. Fig. 1B). Among these classified contigs, 83% were phages, and 13% eukaryotic viruses (Suppl. Fig. 1C), and the remaining contigs were bacterial or plasmid DNA. Most phages belonged to the *Petitvirales* order (52% of phages), which is consistent with previous research^46^, and the most common viruses were from the *Smacoviridae* family (31 % of viruses, Suppl. Fig. 1D).

An overview of the abundance matrix of these 2480 most abundant contigs, grouped by clades, is shown Fig. 1 (see full matrix in Suppl. Table 5). It should be noted that ssDNA viruses abundances cannot be compared to dsDNA ones (*Caudovirales*), due to their over-amplification during the MDA step ^47,48^. Still, *Petitvirales* represented 30% of total abundance, followed by non-bacterial viruses (21%) and *Tubulavirales* (7%). *Tubulavirales* contigs (described in detail below) were grouped according to their type of morphogenesis protein, encoded by *gpI*. We noted a gradient of abundance in *Tubulavirales*, with adults containing most of them (samples are organized in the matrix following this gradient). Among *Caudovirales*, some of the contigs were grouped into higher order taxons (see below). In these cases, the contigs were labelled with the name of a similar reference phage genome when available, and were otherwise assigned a vCONTACT2^36^ cluster number (followed by the size of the circular or largest contig). Still, most signals came from unclustered contigs. We noted that piglets had significantly higher abundances of *Caudovirales* compared to adults (P=0.0036, Student t-test), and fewer ssDNA eukaryotic viruses (not significant, P=0.07). The remaining lower abundance contigs were also classified (see Materials and Methods) and 72% were found to be viral. We conclude that the majority of contigs from these highly purified viromes were indeed viral in nature, with a substantial proportion being ssDNA viruses.

**Figure 1:**
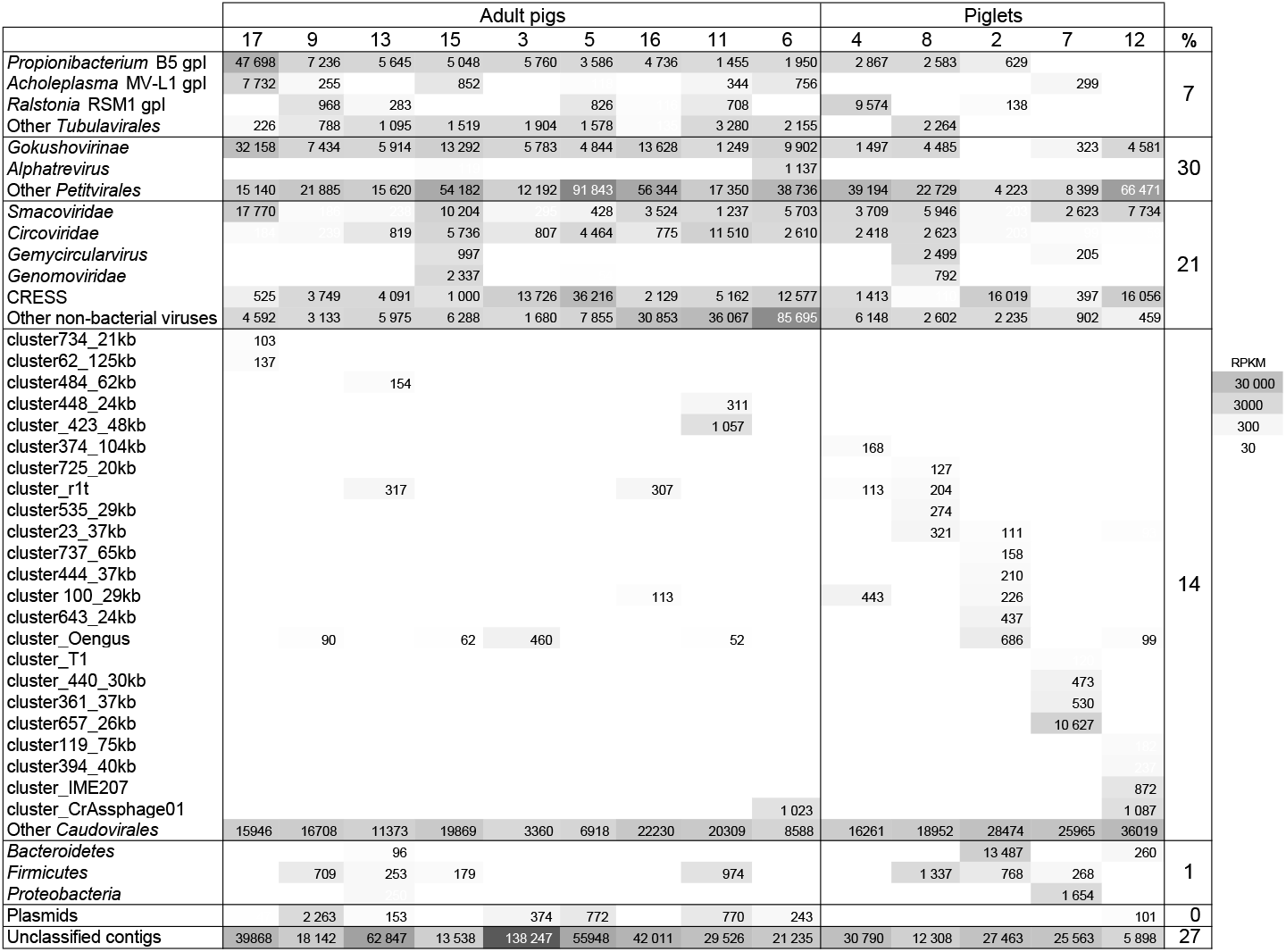
Abundance matrix of the 2480 most abundant virome contigs, grouped into clades. The last column shows the percentage of total RPKM in each contig category: *Tubulavirales, Petitvirales*, non-bacterial viruses, *Caudovirales*, Bacteria, Plasmids, and Unclassified.

### The few antibiotic resistance genes found in viromes are not encoded by phages

ARG were then searched among the complete set of 8090 VP contigs of a size above 2 kb (before clustering, and regardless of their abundance) in viromes. Contigs that were not predicted to be viral were also included (Table 1, see Materials and Methods).The search against ResFam profiles^33^ revealed 8 VP contigs encoding 10 putative ARG, 5 of which were also retrieved with a BLASTp search against MUSTARD^34^. The 5 missing ARG encoded ABC transporters, a category which is simply not included in MUSTARD. Finally, a more stringent search of confirmed ARG with Resfinder^35^ reported only 3 of the 10 initially uncovered ones (Table 1). Interestingly, in one of the 8 ARG-positive contigs (VP7_NODE302), a *tet(40)* gene was found that was undetected with the two previous tools.

**Table 1:**
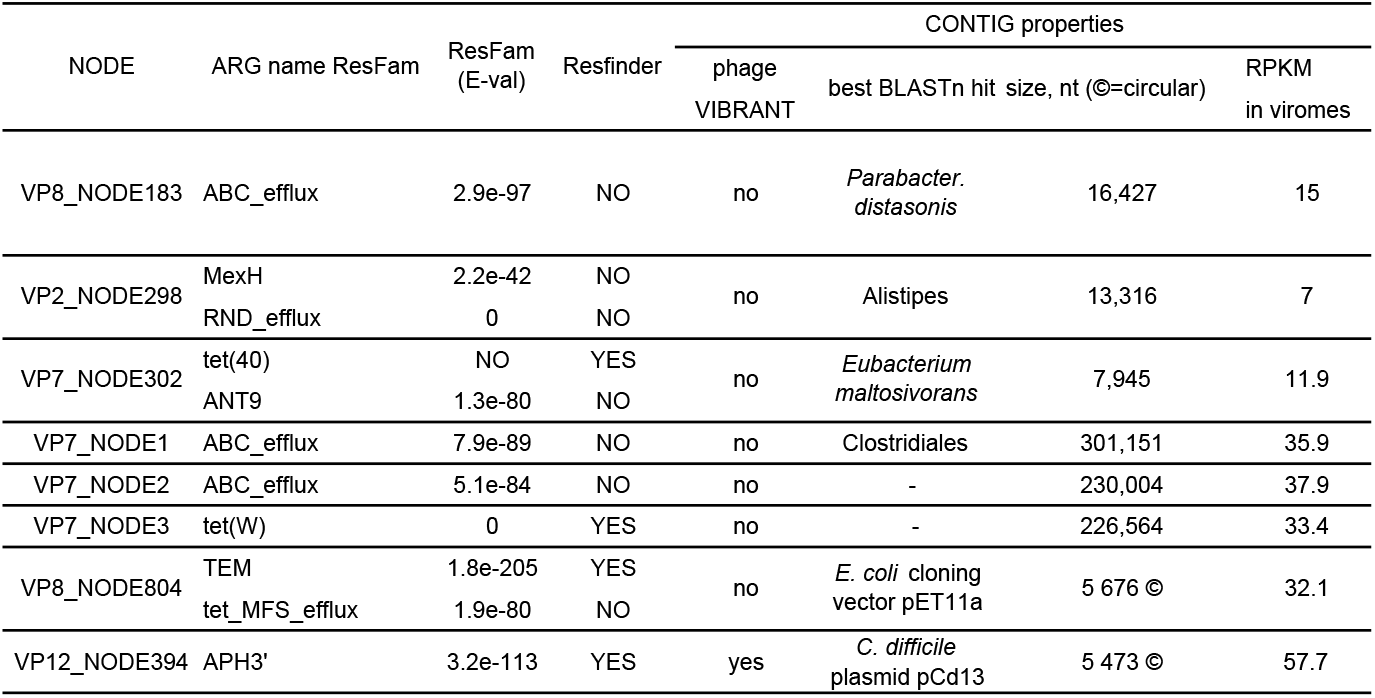
The eight ARG-positive contigs assembled from viromes

Detailed inspection of the 8 identified contigs revealed that none of them originated from phages, despite a positive VIBRANT^31^ prediction for one of them (contig VP12_NODE394, last line in Table 1). This phage candidate, however, turned out to be 99.8% identical to plasmid pCd13 of *Clostridioides difficile* (Genbank accession MH229772). In addition, this 5.4 kb contig was circular and contained a 1.5 kb deletion compared to pCd13. We therefore conclude that it is rather a plasmid than a phage. Of note, the second circular contig, VP8_NODE804, was 100% identical to synthetic plasmid pET11a with its *Nde*I restriction site mutated. We think this laboratory vector comes from the control DNA of the Genomiphi kit used to prepare DNA prior sequencing, suggesting that the sequencing depth was sufficient to detect even minor contaminants. Homologs for all remaining, non phage contigs were searched with BLASTn against NCBI nt database (May 2019) and most of them had a taxonomic association that is typical of intestinal bacterial species (Table 1).

We noted that all ARG-positive chromosomal contigs came from the viromes with less viral enrichment (Suppl. Table 2). Given the lower level of purity, it seems likely that these fragments came from remaining free bacterial DNA rather than viral particles. In contrast, plasmid pCd13 from *Clostridioides difficile* came from a highly purified virome, making bacterial contamination unlikely. This plasmid may have been carried over by generalized transduction.

Taken altogether, the few ARG-positive contigs we found in highly purified pig viromes were not phage-encoded, even in samples originating from farms with heavy antibiotic use, such as sample 16. Compared to the 5806 contigs positively recognized as viral, this places the frequency of ARG-encoded viral contigs below 0.02%. The maximal ARG gene ratio over total viral orfs was below 1.2 10^−5^, or below 0.015 per Mb of assembled viral contigs.

### Pig viromes are rich in inoviruses and Oengus-like phages

Following the examination of ARGs, we explored the main characteristics of these virome assemblies. Among the rich viral content, two main clades emerged. First, 344 VP contigs (272 of them circular) were found to correspond to inoviruses — filamentous phages performing chronic bacterial infections, during which virions are secreted from the bacterial host without killing it^49^. Inoviruses encode a characteristic morphogenesis protein I (gene name *gpI*) which binds to the bacterial membrane and sets the secretion process into motion. Taxonomically, the best characterized family of inoviruses, *Inoviridae*, is now included into a *Tubulavirales* order and the richness of this order has recently been put to light^29^. The role of inoviruses in bacterial pathogenicity has been recognized in many species (for a recent review see^49^), and even a direct effect on human immunity was reported recently^50^. To our knowledge, the presence of functional inoviruses in intestinal microbiota has not yet been reported.

In the abundance matrix of Fig. 1, inovirus contigs were grouped according to their *gpI* type. Most of the contigs (181 of the 344, 52%) possessed the *gpI* type of Propionibacterium phage B5, the only known filamentous phage infecting a Gram-positive host ^51^. Adult pig viromes were particularly rich in this group, compared to piglets. The 272 complete circular genomes were taken for a deeper analysis (Fig. 2, details about all circular genomes are reported in Suppl. Table 3). Their clustering using vCONTACT2^36^ indicated that they were very diverse, forming 50 clusters of a quality above 0.5, and leaving aside 93 singletons. None of the clusters included any of the 10 reference inoviruses available, not even phage B5. The 118 circular and clustered genomes with a B5 *gpI* are highlighted in yellow on the vCONTACT2 map (Fig. 2A).

**Figure 2:**
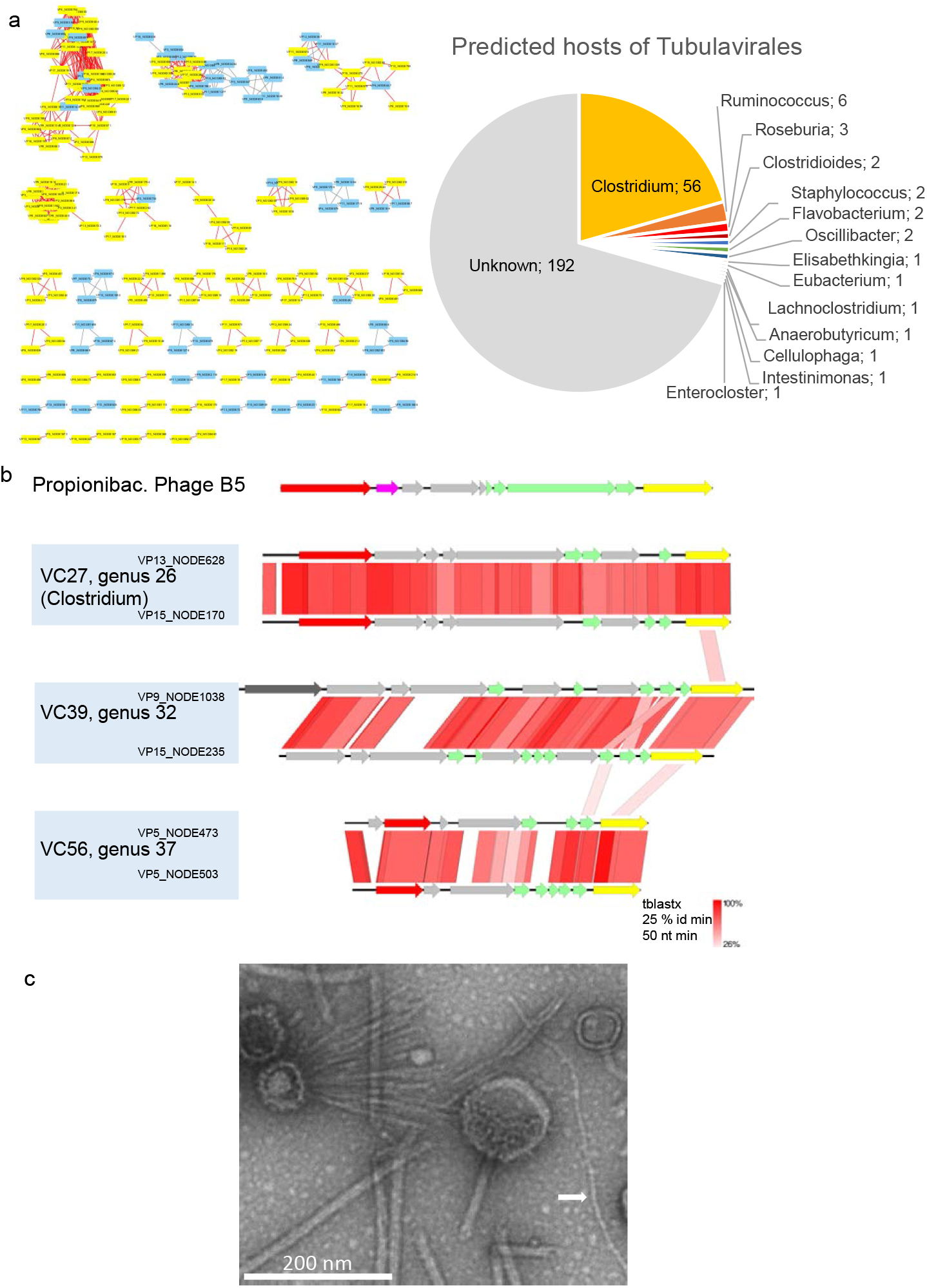
Properties of circular pig inoviruses. **A, left**: VContact2 map of all complete genomes that could be clustered. Those encoding a *gpI* homologous to the one of Propionibacterium phage B5 are highlighted in yellow; **A, right**, host predicted for all circular genomes. **B**. Genome alignments (tBLASTx) and Easyfig map of several inoviruses encoding a B5-like gpI gene. Yellow: gene encoding morphogenesis protein I, red: rep gene for replication initiation, green: structural genes (with transmembrane domains), pink: ssb (single-strand DNA binding) gene. Grey: gene of unknown function. Dark grey: putative phosphoadenosine phosphosulfate reductase. **C** Transmission electron microscope image of virome VP17, the white arrows point to a thin filament that may correspond to an inovirus (length> 420 nm, width, 8 nm).

Interestingly, the GC content in the pig inoviruses ranged from 19.6% to 52.5%, and therefore below the high GC content found in the *Actinobacteria* phylum, such as for *Propionibacterium freudenreichii* (67% GC), the host of the phage B5. We tried to predict the bacterial hosts of these inoviruses with circular genomes. A search of nearly-identical CRISPR spacers led to the prediction of four possible Firmicute hosts: two *Roseburia*, a *Ruminococcus* and a *Lachnoclostridium*. A complementary search using WiSH^45^ or BLASTn added 67 predictions, most often in the *Clostridium* genus (Fig. 2A, right). Consistent with these predictions, a filamentous phage infecting *Clostridium acetobutylicum* NCIB6444 was isolated (but its genome not sequenced) 30 years ago^52^.

Pairwise tBLASTx genome comparisons between B5 and some of the VP genomes with a similar *gpI* revealed similar genetic organizations, although the B5-encoded *gpI* was too distantly related to those of pig genomes to appear in the results (Fig. 2B). Finally, we noticed that electron transmission microscopy images of the VP17 virome, which contained the most inoviruses, revealed some long filaments that are typical of inoviruses (Fig. 2C). We conclude that novel, Firmicute-infecting inoviruses are present in pig fecal samples.

The second notable characteristic of these pig viromes was identified by performing a vCONTACT2 clustering analysis^36^ of the 2991 *Caudovirales* VP contigs, together with 2214 reference phage genomes (see Materials and Methods). Among the seven most populated clusters (listed in Suppl. Table 4), there was a particularly large one of 98 elements, 94 of which originated from pig virome contigs. Of note, four reference phage genomes were connected to them. Three of these four phages are virulent and infect *Actinobacteria*, including the Rhodococcus phages ReqiPoco6 and ReqiPepy6^53^, and Arthrobacter phage Mudcat (accession NC_031224). The fourth is the temperate phage Oengus, which infects the *Firmicute Faecalibacterium prausnitzii*^38^.

The 17 largest genomes of this cluster (45-62 kb) were then compared to already classified phage genomes with ViPTree^54^ (Fig. 3A). The ViPTree branch lengths suggest that the vCONTACT2 cluster includes more than one viral genus. ViPTree also shows that compared to *Actinobacteria* phages, phage Oengus is more closely related to all but one of these VP contigs. Therefore, we named this cluster ‘Oengus-like’. Surprisingly, among the 38 genomes that were larger than 40 kb, only three contained an integrase. This suggests that this clade contains a mix of temperate and virulent phages, with perhaps a small fraction of temperate phages and a majority of virulent phages, at least in the pig viromes. Using Oengus as a reference, whole genome alignment (tBLASTx) was performed for two complete genomes, and it showed that only a few genes were shared among phages of this cluster, confirming further that this cluster represents a taxon beyond the genus level (Fig. 3B). Finally, the search for predicted hosts was performed. The most frequently predicted hosts belonged to the *Clostridiaceae* family (Fig. 3C). In addition, for two of them, a few matching CRISPR spacers were found in strains of *Anaeromassilibacillus*, a genus that belongs to the same *Oscillospiraceae* family as *F. prausnitzii*, the host of phage Oengus.

**Figure 3.**
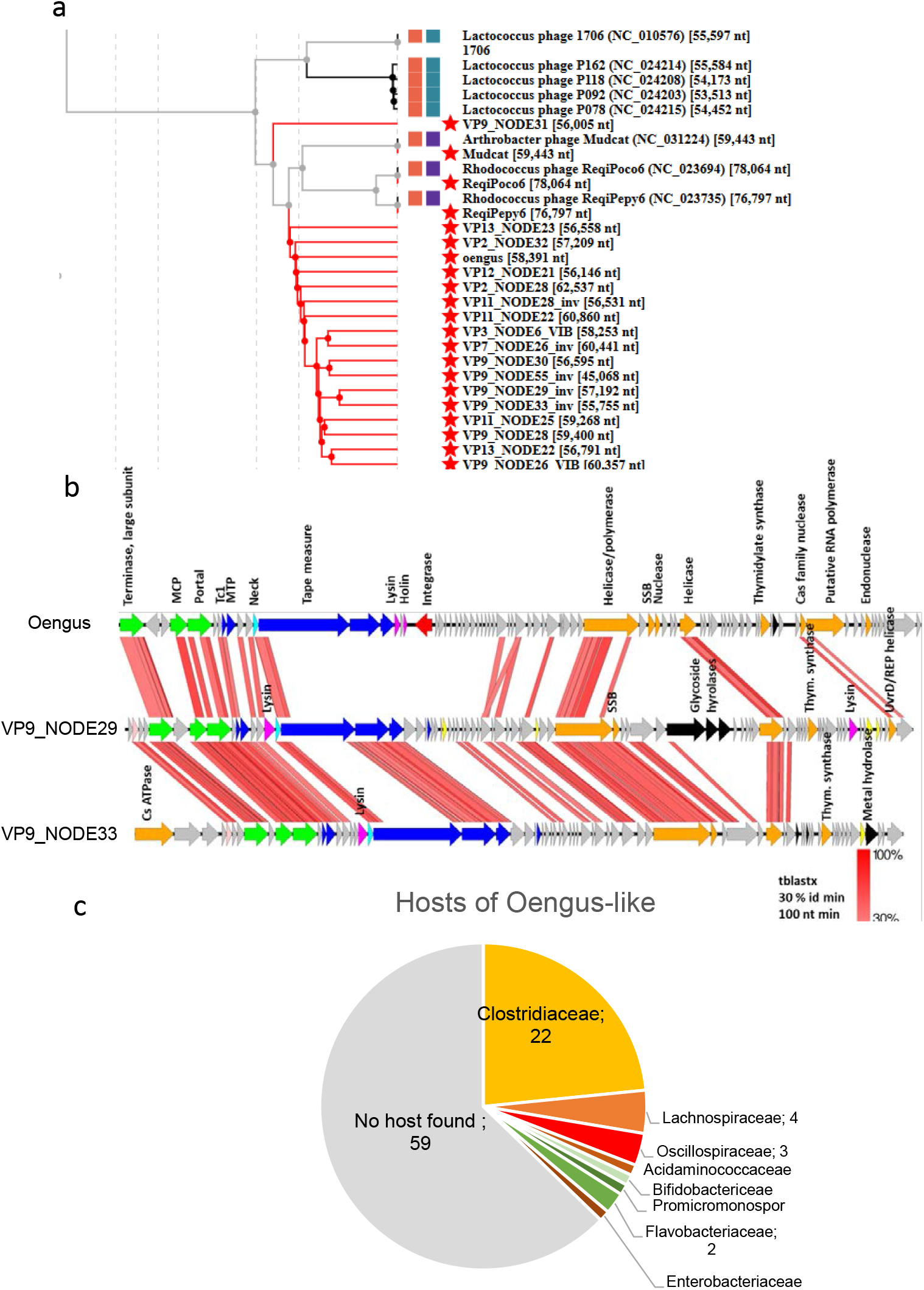
Characteristics of the Oengus cluster. A. Tree representation of the 17 largest Oengus-like contigs together with close reference phage neighbors in the global VipTree^54^, which builds a distance based on shared genes, starting from nt sequences and tBLASTx comparisons. Red stars indicate all genomes that were added to the reference VipTree. **B**. Whole genome alignments (Easyfig map, with tBLASTx option) comparing two relatively close VP nodes (see A) and Oengus. MCP: major capsid protein, Tc1: Tail completion 1, SSB: single-strand binding protein **C**. Predictions of infecting hosts for the 94 contigs of the Oengus-like cluster.

### Pig microbiota contains a high abundance of viruses in addition to six main bacterial phyla

We next investigated whether ARGs could be detected on prophages that are embedded in bacterial genomes. For this, we extracted, sequenced and assembled the DNA from the global fecal samples, following a distinct protocol for DNA extraction (see Materials and Methods, and Suppl. Fig. 2 for the overall analysis pipeline). After assembly and dereplication, a final binning step was performed to aggregate contigs from the same microbial species, using the VAMB tool^26^. To estimate the main phyla present in the 14 pig fecal samples, a taxonomic affiliation with Kaiju was performed on the 220 000 contigs of a size above 2 kb.

In addition, we searched systematically for viral contigs among this microbial contig collection, using a combination of 3 criteria: (i) positive viral result according to VIBRANT^31^, (ii) homologous to a viral VP contig according to BLASTn results, (iii) viral affiliation by Kaiju (see details in Materials and Methods). Of the 220 000 contigs, 16 940 (7.7 %) were deemed of viral origin (labelled viral-P contigs below). We reasoned however, that some of these contigs might correspond to uninduced prophages and some may even be defective prophages. We therefore used the collection of virome reads to determine which of the viral-P contigs were indeed functional (i.e., forming viral particles). Among these 16 940 viral-P contigs, 23.7% were covered by a significant amount of virome reads and classified as “active phages” (see Materials and Methods and Suppl. Fig. 2).

Figure 4 reports a synthetic representation of the main phyla detected, among OTU with abundance above 10 RPKM across the 14 samples (see details in Suppl. Table 6). In this matrix, active phages were binned and counted separately (798 contigs, binned into 547 “active phages”). The remaining viral contigs, for which no proof of activity was obtained, were either kept under their bacterial host taxonomy when available (329 contigs), or placed under “inactive prophages” otherwise.

**Figure 4.**
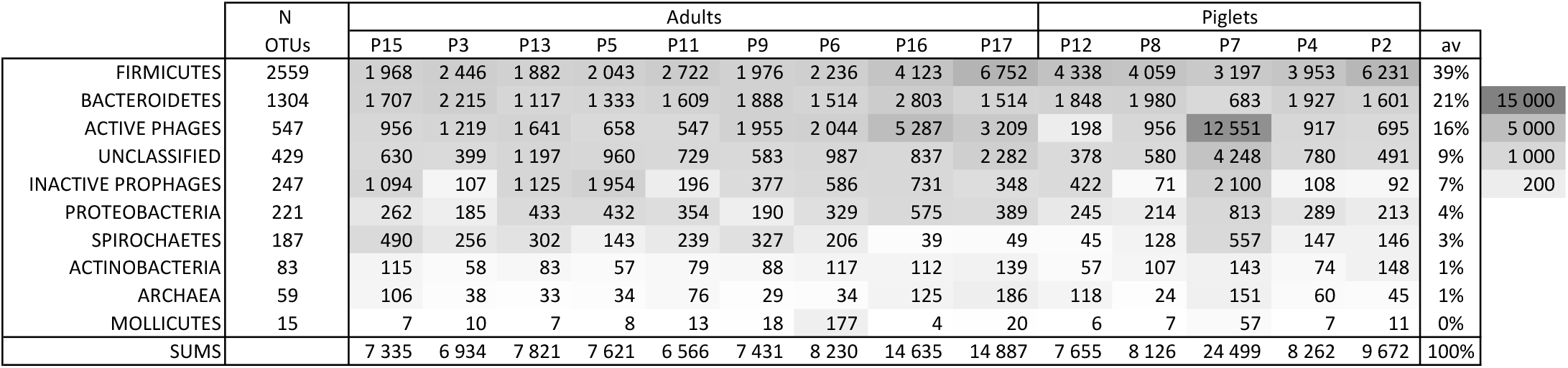
Main pig microbiota phyla. Abundances are expressed as RPKM (grey scale on the right). The average column does not include sample P7

In most samples (13/14), the three main phyla were *Firmicutes* (39%), *Bacteroidetes* (21%) and surprisingly, active phages (16%). The next bacterial core phyla with abundances above 1% were *Proteobacteria*, *Spirochaetes*, *Actinobacteria* and Archaea. Aside from the active phage category, previous reports on pig microbiota described similar phyla distributions ^1,55,56^. The *Mollicute* class was added in the matrix to highlight sample P6, which contained a particularly high level of a single *Mycoplasma* OTU, suggesting pig 6 was sick.

The last sample (piglet P7) had a distinct profile, as virus reads represented 51% of the total. More precisely, a bin of two contigs (12.6 kb total) was responsible for 52% of all viral signal. Upon closer inspection, these contigs had a typical *Picovirinae/Raketienvirinae* organization (Suppl. Fig. 4), but its host could not be predicted. Interestingly, the P7 sample also contained a very large 14 Mb OTU (401 contigs). A BLASTn search against the nt database of the NCBI indicated that its closest (77% identity) and most frequent (18% of contigs from this bin) parent was *Blastocystis hominis* (Eukaryote, Stramenopiles). We also noted that in this sample, richness and abundance in *Bacteroides* and *Prevotella* species was low. This suggests that there is intense predation on bacteria in the microbiota from sample P7.

### A small fraction of ARGs are found in prophages in pig microbiota

An ARG search among the 271 163 P contigs (before dereplication) retrieved 1764 ARG with ResFam and 279 with Resfinder (see Suppl. Table 7 for overall counts, and Suppl. Table 8 for the details of each contig). The large increase in ARG with ResFam is due to the inclusion of ABC transporters, which probably include false positives. We therefore concentrated on ResFinder numbers. The Resfinder counts correspond globally to 1.8 (+/-0.5) 10^−4^ ARG gene per total genes, and 0.18 (+/- 0.05) ARG gene per Mb.

To further investigate the samples with the highest abundances of ARG-positive bacterial contigs and the category of antibiotic resistance that was prevalent, we focused on the abundance matrix of the 168 contigs that harbored the 279 ARG genes (Suppl. Table 8). Overall, a four-fold difference was observed between the most and least ARG-abundant microbiota samples (Suppl. Fig. 5). Piglets also had higher prevalence of ARGs compared to adults. Contigs encoding tetracycline resistance genes were by far the most abundant, followed by aminoglycoside and macrolide resistance genes (Suppl. Fig. 3), consistent with previous reports^1^. Among the *tetR*-encoding contigs, three were particularly abundant, in all samples: the most abundant one encodes a *tet(W)* gene (in P4_NODE_19877) which apparently belongs to an integrative conjugative element (ICE, see Materials and Methods). The two next most abundant contigs (*tet (40)* genes in P12_NODE_17183, 2004 bp, and P8_NODE_13797, 2606 bp) were partially overlapping and matched a plasmid reported from a study on the tetracycline resistome of bio pig farms^2^. Clearly, and as concluded also by Kazimierczak et al., even though tetracycline which was used as a growth factor, is no longer used as such in Europe, the previously selected resistance genes still remain in pig microbiota. Some farms, however, and particularly farm 6, were markedly ARG-poor. What parameter of the farm settings could explain this good record is not known at present. We conclude that globally, the microbial ARG content of these pig fecal samples was similar to that previously reported in the literature^1,57^.

We next sought whether some of the ARG-positive contigs were viral. Three (of 16 940 viral-P contigs, 0.02%) were viral, and all three belonged to the inactive prophages subset (Fig. 5A). These three contigs represent 1.7% of all 168 ARG-positive contigs, while 30 at least were present on other mobile genetic elements (plasmids and ICE, as determined by a BLASTn search against nt, see Suppl. Table 8). A genetic analysis of the 3 ARG+ viral contigs revealed an ICE-prophage hybrid in one case, an IS just next to the prophage region in the second case, and a possible phage gene interruption by the ARG in the last case (star above the gene, Fig. 5B). All together, we conclude that our search for ARG-positive viral contigs across viromes and microbiomes retrieved none in the viromes, and 3 on inactive prophages.

**Figure 5:**
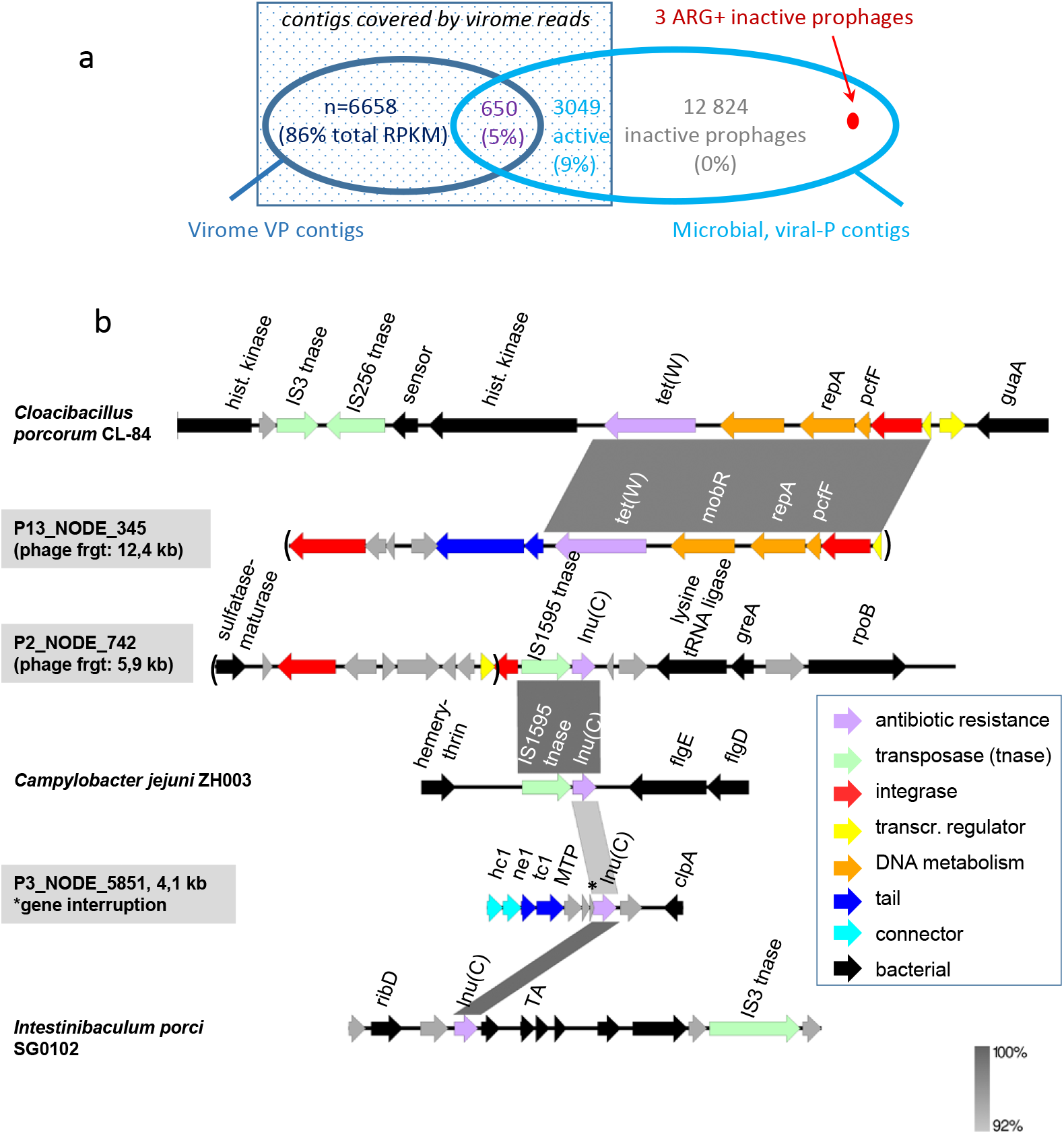
Overview of the ARG found in pig phages and prophages contigs. **A**. Overlap between viral contigs assembled from virome fractions (VP) and those recognized as viral within microbiota assemblies (viral-P). Among 23 181 dereplicated viral contigs (see Materials and Methods for details), 650 were present in both contig sets, while 6658 were present exclusively in viromes. No ARG was found in these two first subsets. Among the viral-P contigs, we distinguish those “covered” by viromes reads, and therefore considered active phages (square of dark blue shade, abundance in viromes indicated as % of RPKM among the 3 subsets of contigs covered by reads), from those that correspond to dormant prophages The 3 ARG positive viral contigs (indicated in red) are among dormant prophages. **B**. Genetic maps of these three prophage regions. Easyfig^42^ maps (BLASTn) of shared regions with other mobile elements are shown. Phage hallmark genes (tail and connector) are colored in blue. GNAT stands for GNAT family N-acetyltransferase, TA for toxin-antitoxin system, GGDEF for GGDEF domain-containing protein.

### Huge phages are detected among microbiota and virome contigs

As a last look at phage contigs present within the microbiota, we attempted to detect any so-called “huge phage” (size > 200 kb)^40^. Indeed, such phage genomes were initially detected in whole microbiota samples, and several were found in pig fecal samples. No close homolog of the previously reported huge phages was found in our dataset, but four distant relatives of complete phage AOT2015-SM01_PERU_trim_clean_scaffold_10 (PERU for short, 201 kb) were found (see Materials and Methods for details). For the first of these, a long contig was present both in the microbiome and virome set. In the microbiome, this contig (P12_NODE_16) was 183 kb long, and its central section matched perfectly with the virome contig VP12_NODE3 (123 kb), confirming its viral nature. The three additional contigs (175-154 kb) came from three viromes VP2, VP9 and VP12. A *Clostridium* host was predicted for one of them (VP2_NODE1) because of a CRISPR spacer^43^. The largest of these 4 contigs was manually annotated, and all 4 were aligned with PERU using tBLASTx and Easyfig (Suppl. Fig. 5). A fraction of the structural module (head genes) and the whole replication module genes were shared between these phages. Interestingly, P12_NODE_16 had long tail genes (absent from PERU and the 3 other contigs) and no detectable sheath encoding gene, suggesting that it is a siphovirus. A few ribosomal genes and eukaryote-like ribonucleoprotein genes (major vault-like protein-encoding genes) were also predicted.

In conclusion, in addition to the two prevalent phage clades found in pig viromes, pig microbiota analyses highlighted a group of four nearly complete phages that are distantly related to a huge phage. A small, highly abundant phage genome with an organization similar to *Picovirinae/Raketienvirinae* was also identified in a dysbiotic sample (P7).

## Discussion

In-depth characterization of 14 pig fecal samples revealed that ARGs are present at 0.18 genes per Mb in the total microbiota of these samples (168 contigs, 279 ARGs). However, none of the ARG-containing contigs appears to belong to active phages, since ARGs were not detected in the viral contigs assembled from viromes reads (detection limit of 0.015 ARGs per Mb of viral contigs). Assuming an average genome size of 3 Mb for bacteria, and 30 kb for phages, these ARG numbers correspond to 0.5 ARG per bacterial genome, and below 3.6 10^−4^ per phage genome. This suggests that ARG genes are at least 1000-fold less likely per viral genome, compared to bacterial genomes. We therefore conclude that even in pigs, and in samples originating from farms with heavy antibiotic usage (such as sample 16), the probability of detecting an ARG on a phage contig is at least 1000-fold lower compared to a bacterial genome.

An explanation for this contrast could be that ARG, whenever they enter prophage genomes (by transposition or recombination), tend to mutate the recipient phage, which loses then rapidly its fitness. However, temperate phage do encode other morons, which probably “landed” as abruptly as ARG in the first place in their genomes. So something specific to the ARG might be at play. For instance, as early as 1947, phage-antibiotic synergy (PAS) was investigated^58,59^ and sub-inhibitory concentrations of antibiotics were shown to be beneficial for phage growth. Recent reports confirmed this effect for virulent phages of *Staphylococcus aureus*^60,61^, *Escherichia coli*^62,63^, and other species^64,65^, although some results suggest the opposite^66^. If this behavior occurs in the gut, phages should not encode genes that diminish this beneficial effect. Importantly, it is unknown whether temperate phages also exhibit this PAS synergy. From the bacterial point of view, one could also argue that an ARG-containing prophage is so beneficial for the host that phage domestication rates speed up in order to retain the ARG.

Indeed, comparison of the 168 ARG-positive contigs to the pool of prophages present in microbiota revealed that three of them were prophages. Among these prophages, we introduced a distinction between those corresponding to functional and active phages, producing virions and therefore covered by virome reads, and inactive prophages. A large majority of these prophages were inactive (12 824, 77% of the total pool of prophage contigs), and the three ARG-positive prophage fragments belonged to these inactive prophages (Fig. 5A). This, together with the genetic map of these fragments (Fig. 5B), suggests they are being domesticated.

In fact, the subset of ‘active phage’ contigs in total microbiota samples entailed more than prophages. DNA from virulent phages, such as the dominant *Picovirinae/Raketienvirinae* of sample P7 (Suppl. Fig. 4) was also present. We can propose that the category of ‘active phages’ also includes the ‘virocell’ fraction, namely the fraction of phage particles replicating in bacteria at sampling time. A recent report suggests that in human microbiota, this fraction represents 25% of all viral contigs detected in total microbiota^67^. Interestingly, in our pig samples, the proportion of ‘active phage’ contigs was similar (23%). This pool of active phages constituted an important fraction of the ecosystem, as it represented in average 16% of genome abundance (RPKM) in each sample (Fig. 4). If phage burst size is around 10 in the intestine (it is estimated at 16 for the Lambda phage in the murine intestine^68^), this proportion of 16% would correspond to a maximum of 1.6% of bacteria hosting an actively replicating phage. In one case (piglet P7 sample), the active phage fraction included 50% of total RPKM. This sample hosted as well a putative Stramenopile, and its bacterial richness was strongly decreased, suggesting dysbiosis.

The comparison between viral contigs found in the virome fraction and the total microbiota is somewhat puzzling, though. Only 1097 of the 7755 virome contigs (14%) have homologs in the total microbiota. Few studies have reported similar comparisons yet, but a similar limited overlap has been reported before by Gregory et al.^69^. Following the virocell reasoning, this might reflect the fraction of viral particles that replicate at time T. Another technical reason for the lack of overlap between microbial and virome viral contigs is certainly phage incomplete assemblies, which may produce different fragments of the same phage in the two subsets. Finally, we cannot exclude that the protocol used to extract total microbial DNA also extracted the DNA present in viral particles. As an important fraction is likely ssDNA, it was not sequenced in the microbiota samples (only 28 of the 2498 ssDNA phage contigs identified in viromes belonged to the overlapping set of 1097 VP contigs). Clearly, more investigations are needed to figure out what fraction of the viral world is captured upon microbiota sequencing, but our results suggest it might be representative, and help figure out phage-bacteria interactions in microbial ecosystems.

This pig sample analysis also revealed the significant progress of all tools designed to sort out viral genomes. We could identify 82% of the 2480 most abundant contigs of the 14 virome samples. Among them, two prominent phage clades were distinguished, inovirus infecting Gram-positive hosts (Fig. 2), and dsDNA phages similar to Faecalibacterium phage Oengus (Fig. 3). Even though viral samples had been purified on gradients, some bacterial and plasmid DNA was still present. We could not determine convincingly whether such DNA corresponded to generalized transduction, or to remaining contamination. Among them, three contigs encoded ARG (according to the Resfinder results), two *tetR* genes on bacterial DNA, both coming from the dysbiotic VP7 piglet sample which was also one of the most contaminated samples, one *kanR* gene on a plasmid (Table 1).

In conclusion, the consistent observations that ARGs are rarely encoded by phages is encouraging for the fight against antibiotic resistance, as well as for phage therapy. These results suggest that this type of mobile genetic element should not pose a threat, except in phages with high levels of generalized transduction. This is also good news in terms of safety for the use of phages to combat recalcitrant infections, as with the exception of those with high transduction levels, phages should not lead to ARG spread.

## Acknowledgements

We would like to acknowledge the Migale bio-informatics facility at INRAE for professional support, and Michi Waygood and Amanda Toperoff from Crayon-bleu for editing service. M.B.D. is recipient of graduate scholarships from the FRQNT and Sentinel North. S.M. holds a T1 Canada Research Chair in Bacteriophages. Q.L.B. and F.T. were recipient of a financial support from an INRAE MEM program.

## Authors contributions

M.B, Q.L.B, J.L, M.D, S.M., E.L.C, J. E, O. Z., contributed to the generation and analysis of data and revised the manuscript, F.T., E.M., C.D., C.A. contributed to the generation of data, M.A.P analyzed data, supervised the work and wrote the manuscript.

## Supplementary tables

Suppl. Table 1: Pig samples, and antibiotic treatments

Suppl. Table 2: Bacterial DNA contamination of virome samples

Suppl. Table 3: Properties of circular inoviruses

Suppl. Table 4: vCONTACT2 VP clusters with largest numbers of contigs

Suppl. Table 5: Abundance matrix of most abundant VP contigs from virome samples

Suppl. Table 6: Abundance table of most abundant microbiota, or P contigs

Suppl. Table 7: ARG detection in microbiota, global statistics

Suppl. Table 8: Detailed properties of all ARG positive contigs from microbiota samples

