## Supplemental figures and tables for "Analysis of viromes and microbiomes from pig fecal samples reveals that phages and prophages are not vectors of antibiotic resistance genes"

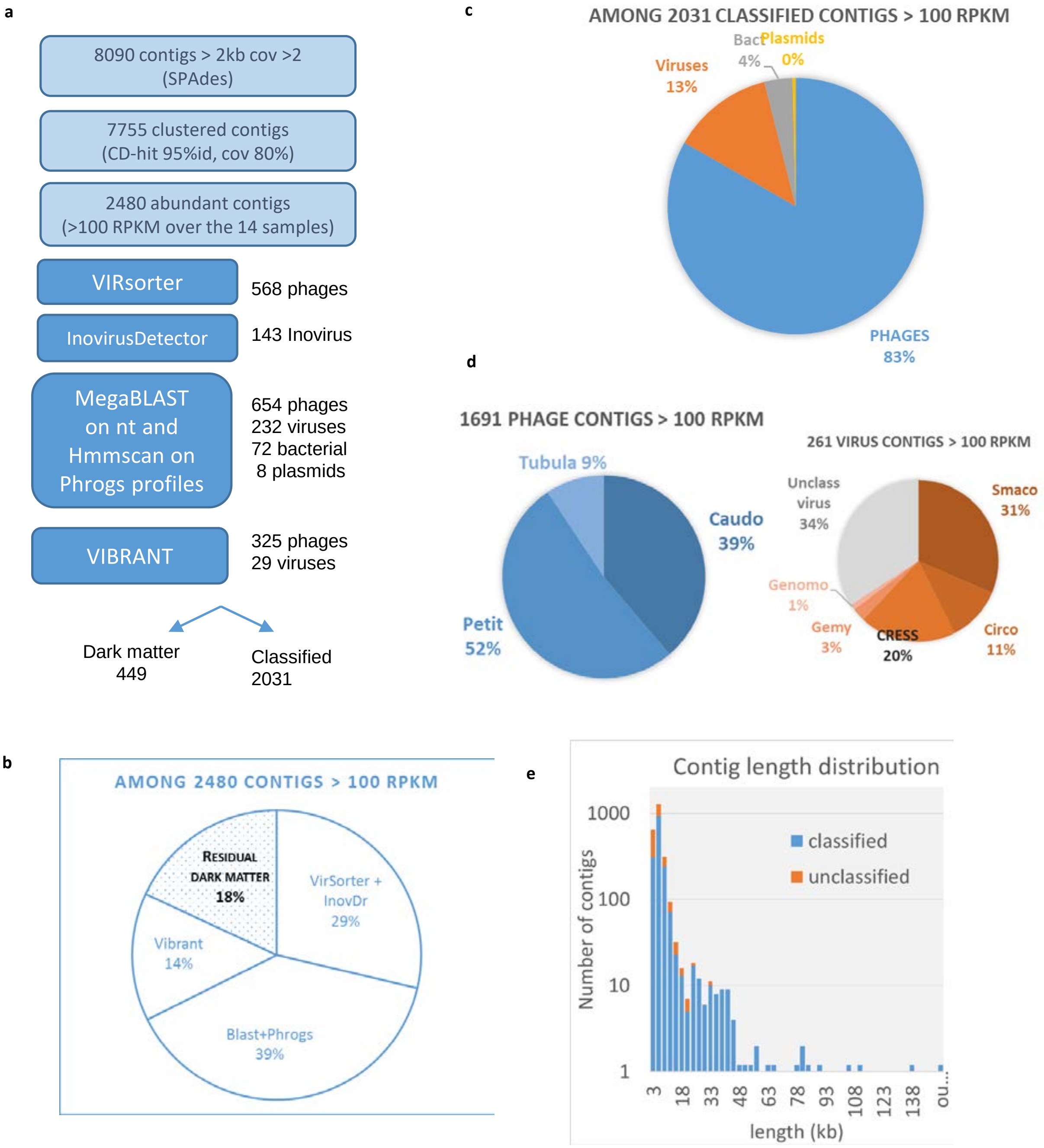


Supplementary Figure 1: Classification of 2480 VP contig with high abundance and size > 2kb. A. Strategy to affiliate contigs. B. 82 % of contigs >2kb are affiliated. C. Most recognized contigs are phages. D. Main categories of phages and viruses. Inoviridae constitute 9 % of all phage contigs, and Smacoviruses is the most populated category of viruses in pig viromes. E. Size distribution of contigs, and proportion of unclassified in each size class.

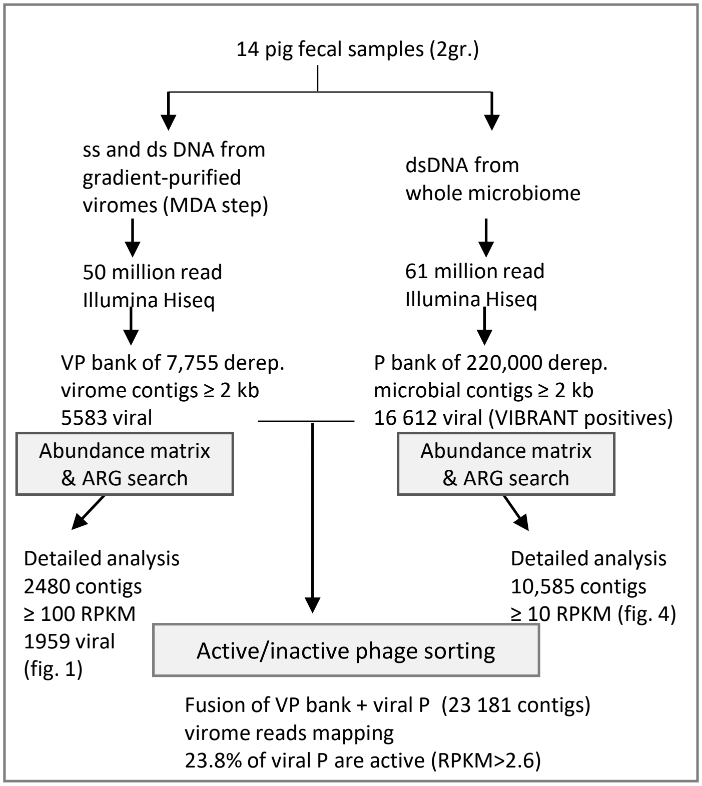


Supplementary Figure 2: Method for detection of viral contigs in microbiomes and partition between active and inactive ones.

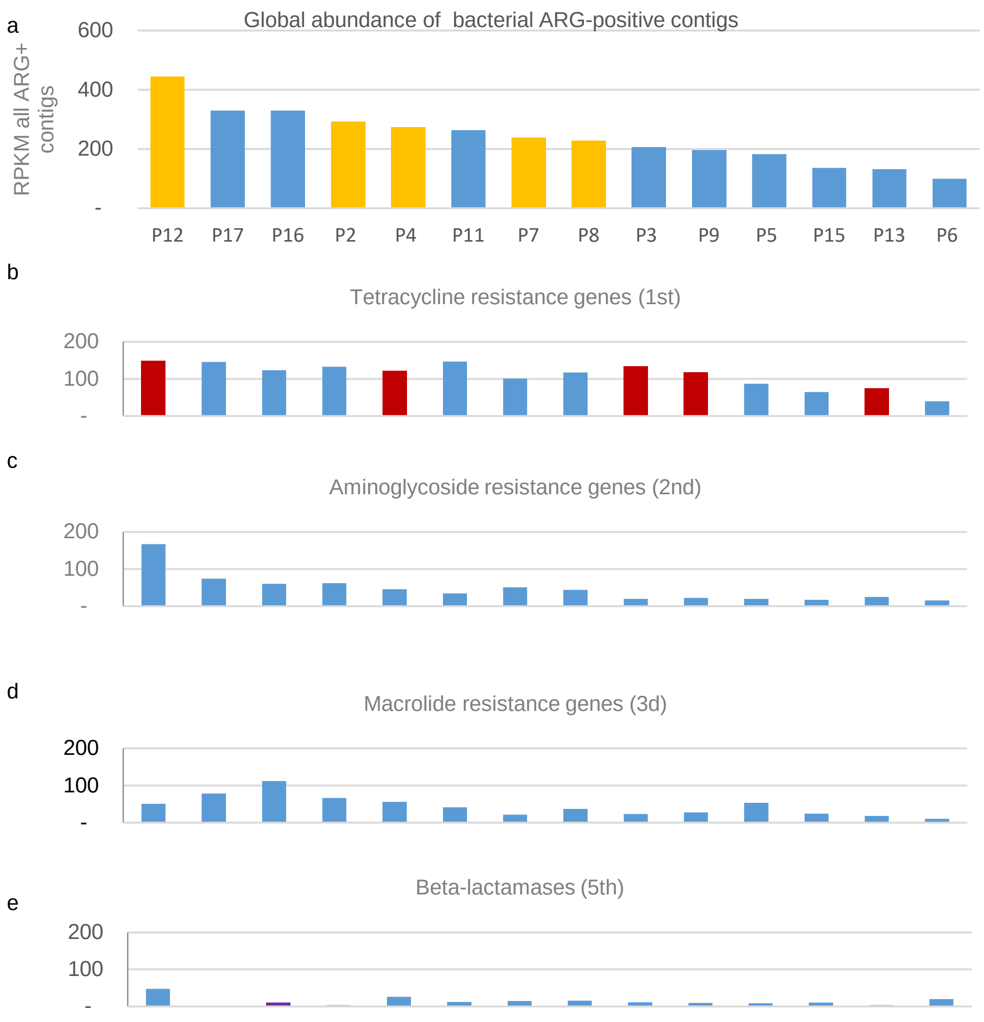


Supplementary Figure 3. **a**. Global abundance of ARG-positive contigs in the 14 pig fecal sample microbiota. Piglet samples are shown in yellow, adults in blue. **b to e**, in the same order as in A, abundance of the main categories of antibiotic resistance genes (rank of abundance. Red bars: animals treated with tetracycline. Purple bar: animal treated with beta-lactams.

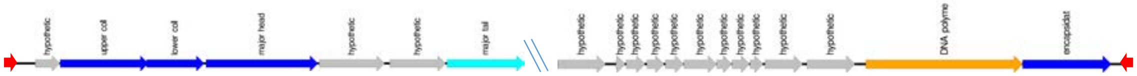


P7_NODE_2713
(5368 bp, 6500 RPKM)

P6_NODE_3950
(7355 bp, 6825 RPKM)

Supplementary Figure 4: The genetic map of the dominant phage in sample P7 reveals a typical Picovirinae organisation. Contigs were annotated with Hhpred, 259 bp terminal inverted repeats are shown in red. The P7_NODE_2713 contig was recognized as viral by VIBRANT, unlike the othercontig.

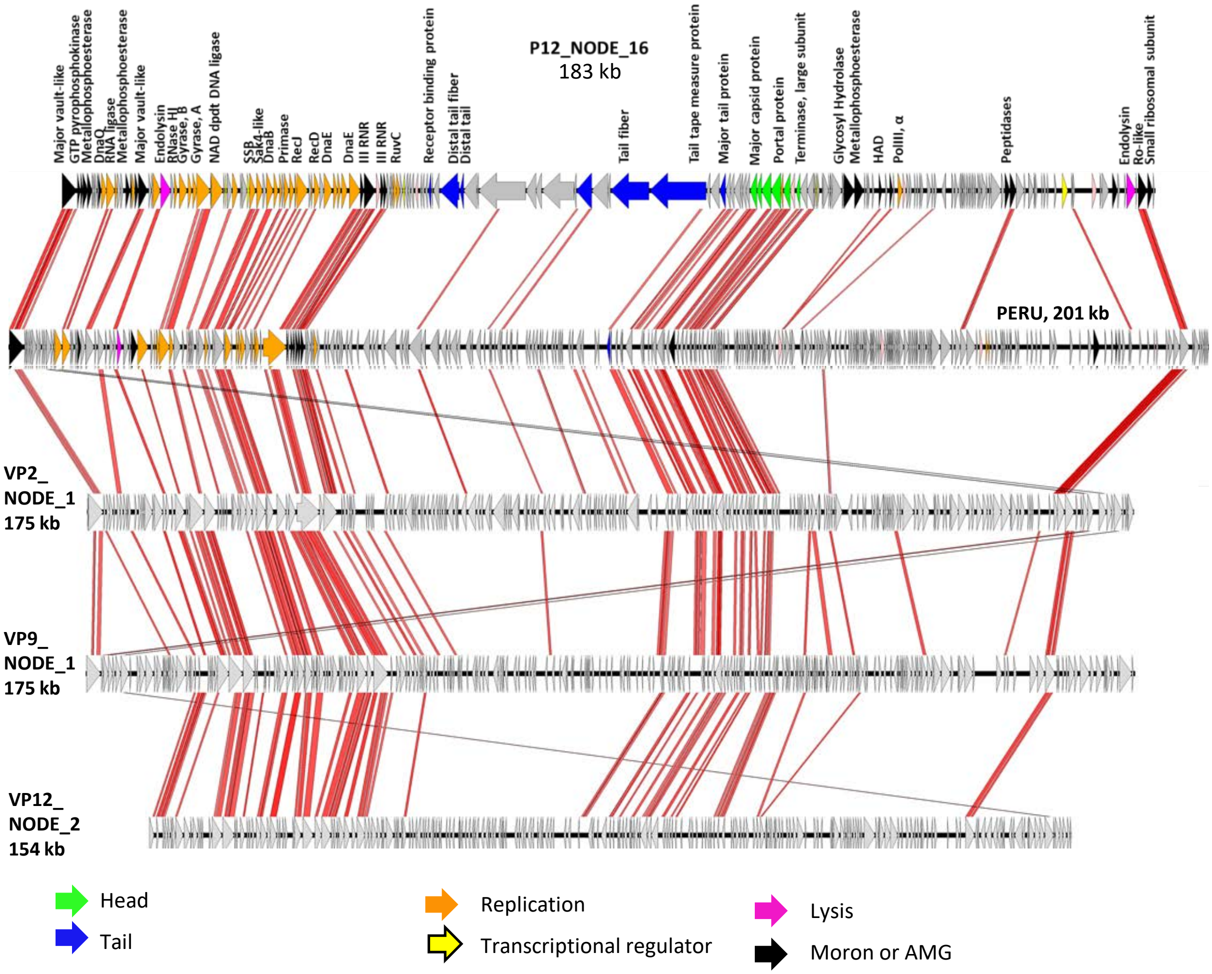


Supplementary Figure 5. Pairwise comparisons between a pig microbiota contig, AOT2015-SM02_PERU (201 kb), and 3 pig virome contigs by tBLASTx, (minimal id. 30%, minimal length 100 bp) using Easyfig. +1 positions were translated on the pig contigs, to fit PERU starting position. P12_NODE6 was annotated manually, gene abbreviations: DnaQ, DnaQ exonuclease; SSB, single- stranded DNA binding protein; Sak4, Sak4-like recombinase; DnaB, DnaB helicase; RecJ, RecJ single-stranded DNA specific exonuclease; RecD, RecD helicase; DnaE, DnaE DNA polymerase III epsilon subunit; III RNR for Class III ribonucleotide reductase; RuvC, RuvC crossover junction endodeoxyribonuclease; HAD, haloacid dehalogenase-like hydrolase, polIII α, DNA polymerase III alpha subunit. P12_NODE16 differs from all other genomes by the presence of clear and long tail genes.

Supplementary Table 1: pig samples, and antibiotic treatments

| sample # | Animal type | Antibiotic treatments on animal sampled |
|---|---|---|
| 3 | adult, female | oxytretracycline |
| 5 | adult, female | |
| 6 | adult, female | |
| 9 | adult, female | oxytretracycline |
| 11 | adult, female | dexalone, marbofloxacine |
| 13 | adult, female | doxycycline |
| 15 | adult, female | |
| 16 | adult, female | TMP, sulfadiazin, tylosin, amoxicillin, marbofloxacin |
| 17 | adult, female | |
| 2 | piglet | millicoli, tiamvet |
| 4 | piglet | oxytetracycline |
| 7 | piglet | |
| 8 | piglet | lyncomycin, colistine |
| 12 | piglet | oxytetracycline |

Supplementary Table 2: bacterial DNA contamination of virome samples

| Sample | Filtered | Total reads | Reads mapped on 16S (Silva) | Freq 16S reads | viromeQC enrichment |
|---|---|---|---|---|---|
| VP2 | | 45 413 231 | 1287 | 2.83E-05 | 7 |
| VP3 | | 38 432 018 | 23 | 5.98E-07 | 100 |
| VP4 | YES | 39 665 645 | 68 | 1.71E-06 | 100 |
| VP5 | YES | 44 131 406 | 2 | 4.53E-08 | 100 |
| VP6 | | 41 894 101 | 44 | 1.05E-06 | 100 |
| VP7 | | 43 350 198 | 769 | 1.77E-05 | 26 |
| VP8 | | 48 854 252 | 281 | 5.75E-06 | 62 |
| VP9 | | 46 636 927 | 171 | 3.67E-06 | 100 |
| VP11 | | 37 192 968 | 104 | 2.80E-06 | 100 |
| VP12 | | 43 095 618 | 63 | 1.46E-06 | 100 |
| VP13 | | 40 804 259 | 104 | 2.55E-06 | 100 |
| VP15 | YES | 39 981 040 | 4 | 1.00E-07 | 100 |
| VP16 | | 43 004 890 | 19 | 4.42E-07 | 100 |
| VP17 | YES | 43 630 846 | 11 | 2.52E-07 | 100 |

Supplementary Table 4: vCONTACT2 VP clusters with largest numbers of contigs

| Name | N VP contigs | ref phage | av size complete genome |
|---|---|---|---|
| cluster_1 | 94 | Faecalibacterium phage Oengus | 58 kb |
| cluster_23 | 38 | None | 38 kb |
| cluster_62 | 17 | None | 125 kb |
| cluster_99 | 17 | Lactococcus phage r1t | 33 kb |
| cluster_100 | 16 | None | 29 kb |
| cluster_88 | 8 | Bacteroides phage B40-8 | 46 kb |
| cluster_328 | 7 | Bacteroides Crassphage01 | 90 kb |

Supplementary Table 7: ARG detection in microbiota, global statistics

| Sample | animal type | N ORF | Mb, all contigs | N ARG ResFam | N ARG Resfinder | ARG ResFam /N ORF | ARG ResFam /Mb | ARG Resfinder /N ORF | ARG Resfinder /Mb |
|---|---|---|---|---|---|---|---|---|---|
| P3 | adult | 98 507 | 97.868 | 102 | 19 | 1.04E-03 | 1.04 | 1.93E-04 | 0.19 |
| P5 | adult | 128 655 | 132.811 | 89 | 26 | 6.92E-04 | 0.67 | 2.02E-04 | 0.20 |
| P6 | adult | 154 692 | 148.622 | 101 | 19 | 6.53E-04 | 0.68 | 1.23E-04 | 0.13 |
| P9 | adult | 143 904 | 141.983 | 98 | 21 | 6.81E-04 | 0.69 | 1.46E-04 | 0.15 |
| P11 | adult | 125 978 | 127.457 | 175 | 26 | 1.39E-03 | 1.37 | 2.06E-04 | 0.20 |
| P13 | adult | 121 095 | 117.966 | 86 | 19 | 7.10E-04 | 0.73 | 1.57E-04 | 0.16 |
| P15 | adult | 124 547 | 120.031 | 100 | 19 | 8.03E-04 | 0.83 | 1.53E-04 | 0.16 |
| P16 | adult | 112 647 | 114.048 | 161 | 31 | 1.43E-03 | 1.41 | 2.75E-04 | 0.27 |
| P17 | adult | 77 636 | 76.678 | 122 | 16 | 1.57E-03 | 1.59 | 2.06E-04 | 0.21 |
| P2 | piglet | 91 912 | 92.012 | 131 | 11 | 1.43E-03 | 1.42 | 1.20E-04 | 0.12 |
| P4 | piglet | 144 329 | 144.556 | 183 | 20 | 1.27E-03 | 1.27 | 1.39E-04 | 0.14 |
| P7 | piglet | 72 106 | 71.658 | 106 | 20 | 1.47E-03 | 1.48 | 2.77E-04 | 0.28 |
| P8 | piglet | 117 983 | 118.835 | 143 | 11 | 1.21E-03 | 1.20 | 9.32E-05 | 0.09 |
| P12 | piglet | 111 326 | 113.541 | 167 | 21 | 1.50E-03 | 1.47 | 1.89E-04 | 0.18 |
| sum | | 1 087 661 | 1 077.46 | 1034 | 196 | | | | |
| average | | | | | | 1.13E-03 | 1.13 | 1.77E-04 | 0.18 |
| std dev | | | | | | 3.54E-04 | 0.35 | 5.49E-05 | 0.05 |
